# Density Fluctuations Yield Distinct Growth and Fitness Effects in Single Bacteria

**DOI:** 10.1101/2021.05.14.444254

**Authors:** Shahla Nemati, Abhyudai Singh, Scott D. Dhuey, Armando McDonald, Daniel M. Weinreich, Andreas. E. Vasdekis

## Abstract

Single-cells grow by increasing their biomass and size. Here, we report that while mass and size accumulation rates of single *Escherichia coli* cells are exponential, their density fluctuates during growth. As such, the rates of mass and size accumulation of a single-cell are generally not the same, but rather cells differentiate into increasing one rate with respect to the other. This differentiation yields a previously unknown density homeostasis mechanism, which we support mathematically. Further, growth differentiation challenges ongoing efforts to predict single-cell reproduction rates (or fitness-levels), through the accumulation rates of size or mass. In contrast, we observe that density fluctuations can predict fitness, with only high fitness individuals existing in the high density fluctuation regime. We detail our imaging approach and the ‘invisible’ microfluidic arrays that critically enabled increased precision and throughput. Biochemical production, infections, and natural communities start from few, growing, cells, thus, underscoring the significance of density-fluctuations when considering non-genetic variability.

## Introduction

Across all domains of life, cell growth relies on a series of biochemical processes through which cells synthesize new components, replicate their genetic material, increase their size and, eventually, divide to increase their abundance^1-4^. As such, growth is a key parameter in cellular physiology^5^, evolution^6^, the production of high-value chemicals^7^, as well as human, animal, and plant health^8, 9^. Recent investigations at the single-cell level have revealed significant variability in the rates of growth among clonal cells^10^. This form of non-genetic variability has been attributed to fluctuations in the abundance of catabolically active enzymes^11^ that generally emanates from the stochastic nature of gene expression^12-15^. Reproduction variability between isogenic cells has also been observed^16-18^. In this context, some cells divide considerably sooner (or later) than the population average, thereby yielding distinct fitness effects that occur at much shorter timescales than what mutations can confer^16-18^.

Commonly, single-cell growth is investigated by recording the dynamics of cell size (i.e., length, area, or volume). These, size-based, investigations have unraveled key size homeostasis mechanisms, including the critical accumulation of division proteins and timing of DNA replication initiation^19-22^. Further, size-based investigations have informed us about mutation dynamics of single-cells and resulting fitness effects^23^, as well as how cellular noise can increase population fitness under steady-state conditions^24, 25^. In parallel, single-cell growth has been also examined by recording their dynamics of mass accumulation^26^. Fundamentally, these measurements capture the underlying metabolic dynamics of nutrient conversion to building blocks, such as amino acids, lipids, and nucleotides^26^. In this context, mass-based investigations have unmasked the exponential nature of mass production^27^, as well as the presence of ATP-driven high frequency mass fluctuations^28^. Moreover, mass-based investigations have revealed that the growth-rate of mammalian cells is not constant across the cell cycle^29, 30^, as well the influence of cellular noise on the metabolic trade-offs between the naturally evolved growth and engineered pathways^31^.

Clearly, growing cells need to coordinate both size and mass accumulation, with the latter being enthalpically more relevant than the former^31^. Cellular size and mass are linked through dry mass density (dry-density henceforth), namely: the number of molecules per unit volume, a metric that represents the level of crowding within the cytosolic environment. Unlike previous, population-level, readouts^32, 33^, single-cell analyses reveal considerable cell-to-cell variability in dry-density, as displayed in **Fig. 1a** with a coefficient variation of 9%; however, and despite the significant discoveries pertaining to cell size regulation^19-22^, the non-genetic variability in density and its influence on the size-mass coordination during growth remain significantly less understood. A notable exception here is a recent report that cellular density varies with the cell’s surface-to-volume ratio, highlighting the importance of biomass growth in cell size control^34^. Importantly, it is also not known how density variability might influence the reproduction rates or fitness levels of single-cells^35, 36^. Indicatively, some reports quantify the fitness of single-cells through the rate of size increase^23^; conversely, mass production rates can also be considered as a fitness metric, given the direct link between the mass production rates of single-cells and the population-level Malthusian parameter^16, 37^. As demonstrated here, however, size and mass accumulation rates are not generally equal, thus, confounding both parameters as predictors of single-cell fitness.

**Figure 1.**
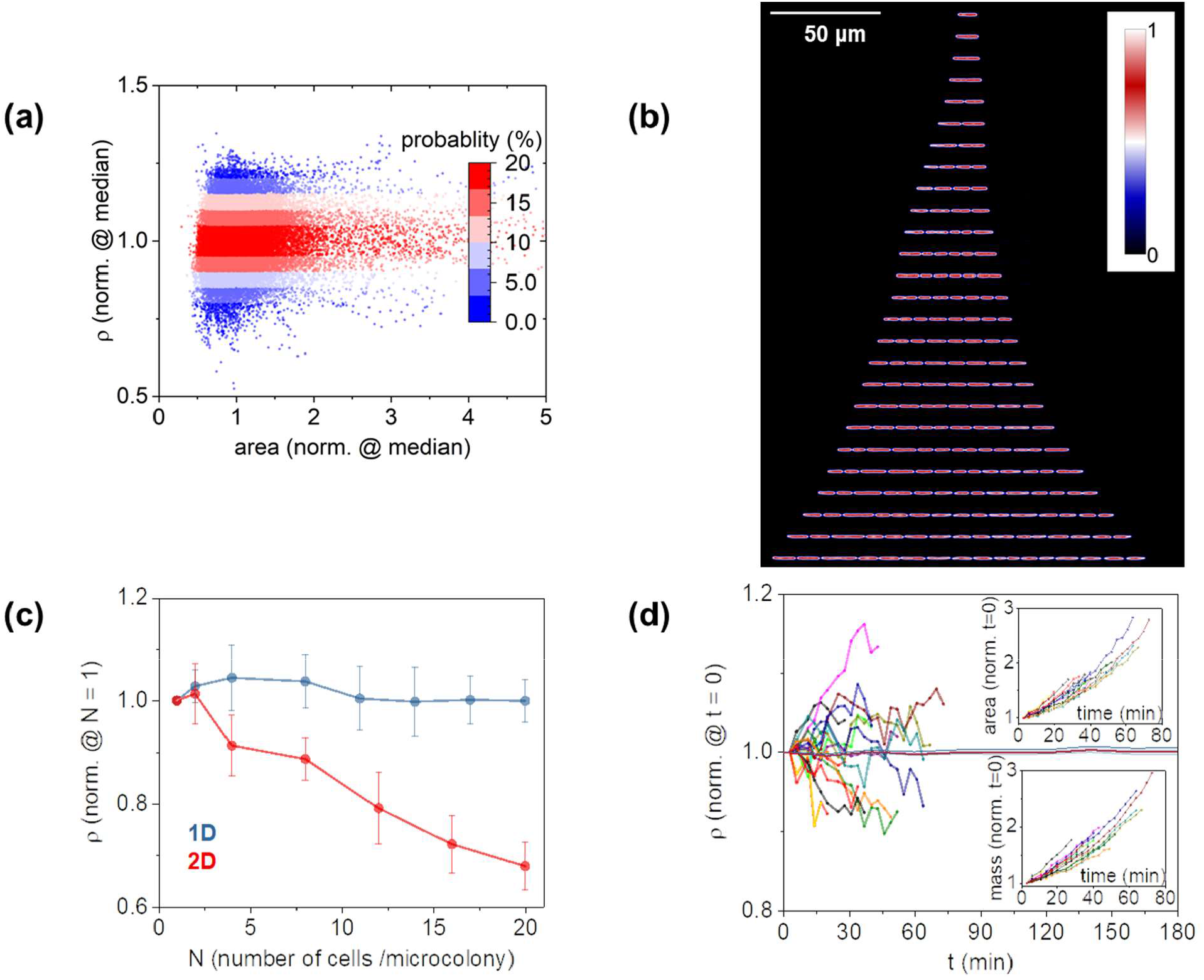
**(a)** Cell-to-cell density variability as a function of cell size (approximated by its area); graph represents the response of single-cell snapshot data at various stages along their cycle (n = 35,000 observations). **(b)** Microcolony expansion from a single-cell to 4 generations using quantitative-mass imaging; the vertical direction in the montage represents time, while color coding represents cell density normalized to the mother density at t = 0. **(c)** Microcolony density (normalized at t = 0) during expansion using 1D and 2D growth assays; data points and error-bars represent the average and standard deviation of n ∼ 8 measurements, respectively; time-dependent density differences in 2D where statistically significant (one-way ANOVA, F(6,49) = 51.8, p << 0.001); no such evidence was found for the 1D assays (one-way ANOVA, F(7,56) = 0.8 and p = 0.5). **(d)** Density, mass, and area growth curves of individual *E. coli* cells from birth to division; all parameters are normalized at t = 0 and color-coding represents the dynamics in size, mass, and density of the same cell; horizontal red line denotes the dynamics of the normalized density of 10 polystyrene particles (1 μm diameter) over time (red line denotes the average and blue-shaded area denotes the 95% confidence intervals).

## Results

### Single-Cell Density Measurements

Addressing these knowledge-gaps requires assays that can quantify the dynamics of both density and size of single, growing cells at high-throughput rates. Quantitative-phase imaging is an ideal candidate to probe these dynamics in a non-invasive manner^30, 31, 38-41^; in these schemes, however, the dynamic nature of microcolony expansion can yield substantial loss of information. Specifically, interferometric imaging schemes that rely on spatially coherent illumination are limited in spatial bandwidth, which can in turn constrain the homogeneity of the reference field (i.e., the halo effect)^42^. Such inhomogeneities become detrimental when multiple cells reside in close proximity (e.g., when growth is confined to 2D^43^), or cells are imaged in the vicinity of dielectric discontinuities (e.g., microfluidic walls). Light scattering between cells^44^, or between cells and dielectric discontinuities^45^ can also incur information loss in modalities relying on spatiotemporally coherent illumination, unless computationally intensive backpropagation algorithms are implemented^46, 47^.

To overcome these shortcomings, we constructed a microarray that enables dynamic density and size tracking of multiple single *Escherichia coli* cells with minimal light scattering between cells and between cells and dielectric discontinuities (**Fig. 1b**). To achieve this, our design relied on two key characteristics. *First*, we adopted an 1D immobilization strategy^48, 49^ that positioned cells at locations that deterministically eliminate cell crowding and the resulting cell-to-cell scattering. *Second*, we employed a polymer matrix that became ‘invisible’ upon contact with water, thus, eliminating scattering between cells and the microfabricated features. Both of these characteristics uniquely enabled the dynamic tracking of single-cell size, mass, and density for up to 6-7 generations (**Fig. 1c**). In this context, nutrients and stimuli were supplied through vertically integrated membranes or microfluidics (**Methods, Sup. Fig. 1**). Further, inspired by Moore’s Law and related strategies in microelectronics, we applied electron-beam lithography to define multiple 1D constrictions at micron-scale distances between them^50^. Critically, this lithography step increased the resulting throughput rates (i.e., the number of observations per unit area) by more than one order of magnitude relative to conventional 2D growth approaches (**Sup. Fig. 2**).

### Growth Differentiation

The combination of the “invisible” 1D microarray with spatial light interferometric imaging revealed that while both size (approximated by cell area, see **Methods**) and mass of single *E. coli* increased exponentially (**Fig. 1d**, *inset*), cellular dry-density was not constant during growth (**Fig. 1d**). Contrary to previous population-level readouts of cellular density^32, 33^, we observed that dry-density undergoes non-monotonic increases or decreases during growth for most cells (**Fig. 1d**). We did not observe such fluctuations in the density of polymer microparticles in similar time-lapse imaging exercises (**Fig. 1d**), suggesting that the observed cell density fluctuations are not due to technical noise. To a similar end, we also observed density fluctuations at higher temporal resolution (**Methods**), characterized by ‘smoother’ variations with time than those displayed in **Fig. 1d**. It is also worth mentioning that density fluctuations have also been reported recently by others in *E. coli*^34^ without, however, any further analyses on these dynamic phenomena. Interestingly, the overall magnitude of these fluctuations varied between cells, while – unexpectedly – some cells exhibited positive and others negative trends on average during their growth cycle (**Fig. 1d**).

We hypothesized that single-cell density fluctuations during growth could be interlinked with the rates of mass or size accumulation throughout the cell cycle (i.e., from birth to division). To assess this hypothesis, we quantified the accumulation rates of size (γ_A_) and mass (γ_M_) during the cell cycle (defined as 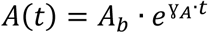 and 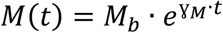, **Methods**). We observed that these two rates were not identical (i.e., γ_A_ ≠ γ_M_), but rather clonal bacteria differentiated into two continuous subpopulations: one that exhibits higher rates of size accumulation (γ_A_ > γ_M_) and one that reverses this behavior (γ_A_ < γ_M_, **Fig. 2a**). As per our original hypothesis, growth differentiation (γ_A_ ≠ γ_M_) was found to be essentially driven by the underlying density fluctuations during the cell cycle. Specifically, cells with density fluctuations that were overall positive during growth differentiated into higher rates of mass accumulation (i.e., γ_A_ < γ_M_); conversely, overall negative fluctuations maximized the rates of size accumulation (i.e., γ_A_ > γ_M_, **Fig. 2b**).

**Figure 2.**
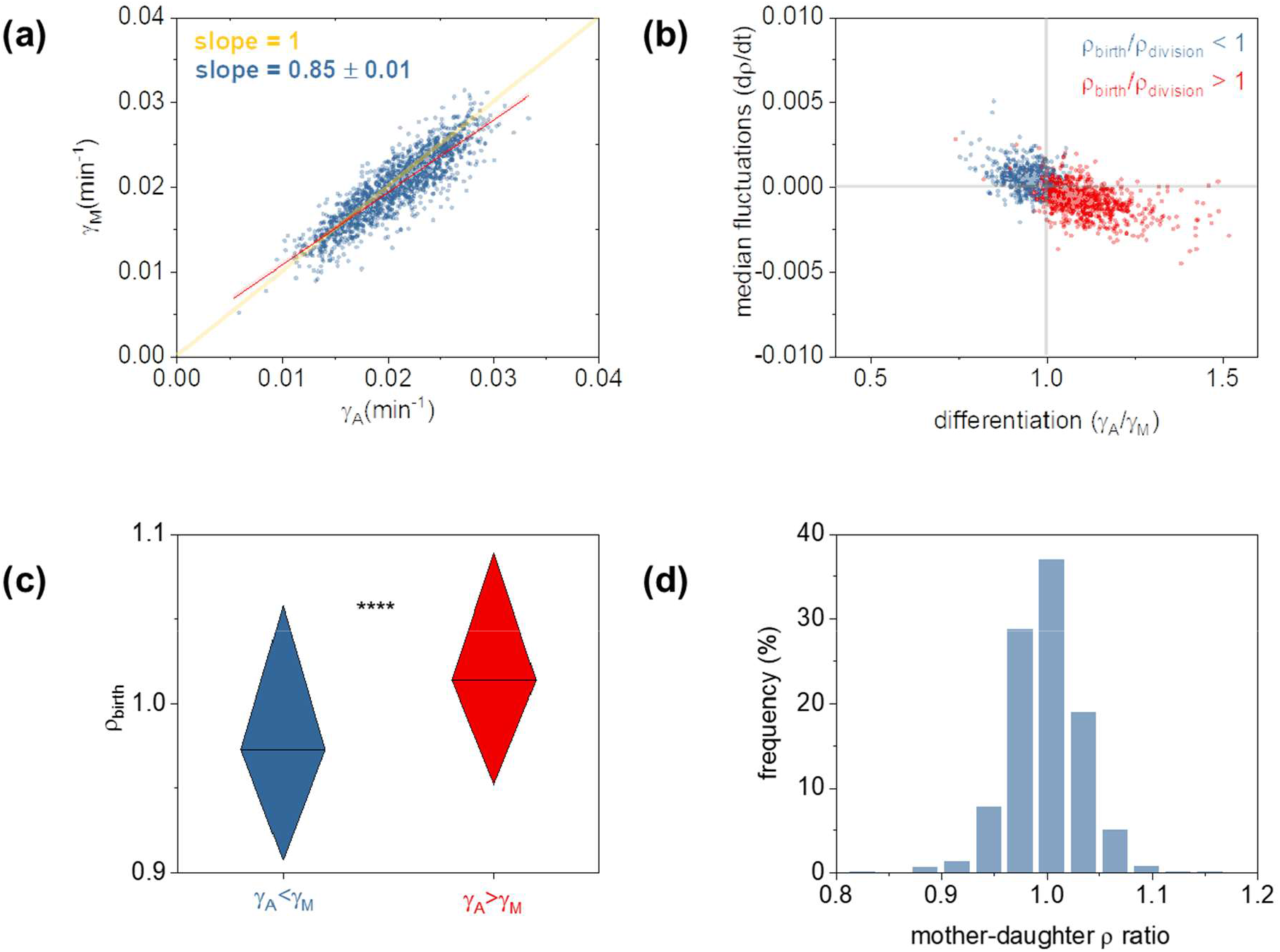
**(a)** Growth differentiation with some cells optimizing area (γ_A_) and others biomass (γ_M_) accumulation rates; graph represents the cumulative response of three replicates, with each replicate presented separately in **Sup. Fig. 3**; red line corresponds the linear fit to the experimental data (shaded areas are the 95% confidence intervals), while yellow line represents a slope of 1. **(b)** Median density fluctuations during growth (dρ/dt, *y-axis*) as a function of growth differentiation (γ_A_/γ_M_, *x-axis*); dρ/dt were calculated as the median value of all density fluctuations (namely: dρ = ρ_i+1_ – ρ in the dt = t_i+1_ – t_i_ timeframes – see *Methods* section) during the cell cycle; similarly, the growth rates in cell mass and size were calculated by exponential fits throughout the cell cycle, as detailed in the *Methods* section; color coding corresponds to increased (blue) or decreased (red) cellular density prior to division; graph plots the cumulative response of three replicates, with each replicate presented separately in **Sup. Fig. 4. (c)** The growth differentiation dependence on the cellular dry-density at birth (normalized over the median); boxcharts represent the 25%-75% of the three replicates combined with each replicate plotted separately in **Sup. Fig. 5**; asterisk denotes statistical significance (Mann-Whitney test: U = 203746, p < 0.001, with additional statistical tests reported in **Sup. Table 1**). In support of this finding, we also plot the differentiation dependence on the density at birth in **Sup. Fig. 6a. (d)** Division asymmetry in dry-density, as noted by the density ratio between mothers (at division) and daughters (at birth). The graph represents the cumulative response of three replicates, with each replicate presented separately in **Sup. Fig. 7**.

In further exploring correlates of growth differentiation, we found that this form of differentiation can be statistically predicted from the cellular dry-density at birth (**Fig. 2c**). Specifically, cells born with lower density than the population median tend to exhibit higher rates of mass accumulation (γ_A_ < γ_M_) and vice versa. Ultimately, density variability at birth can be attributed to the innate stochasticity in cellular physiology. This form of stochasticity can potentially include the asymmetric partitioning of biomolecules upon division. Such asymmetric partitioning was recently evidenced in the division of single gene products between daughter *E. coli* cells^51^. A similar form of asymmetry is also evident in our work **Fig. 2d**).

### Density Homeostasis

Concomitantly, we observed that density fluctuations (and the resulting growth differentiation) decreased under the MIC-level pressure of ampicillin (**Fig. 3a**), a bactericidal antibiotic that inhibits cell wall biosynthesis and division. Specifically, upon exposure to antibiotics, *E. coli* cells exhibited variable phenotypic responses: approximately 12% of the population either died rapidly or stopped growing, while 88% expressed a filamentous, non-dividing phenotype^52^. The observation of decreased density fluctuations pertain specifically to non-dividing filamentous cells and further supports the notion of a potential relationship between density fluctuations and cell division.

**Figure 3.**
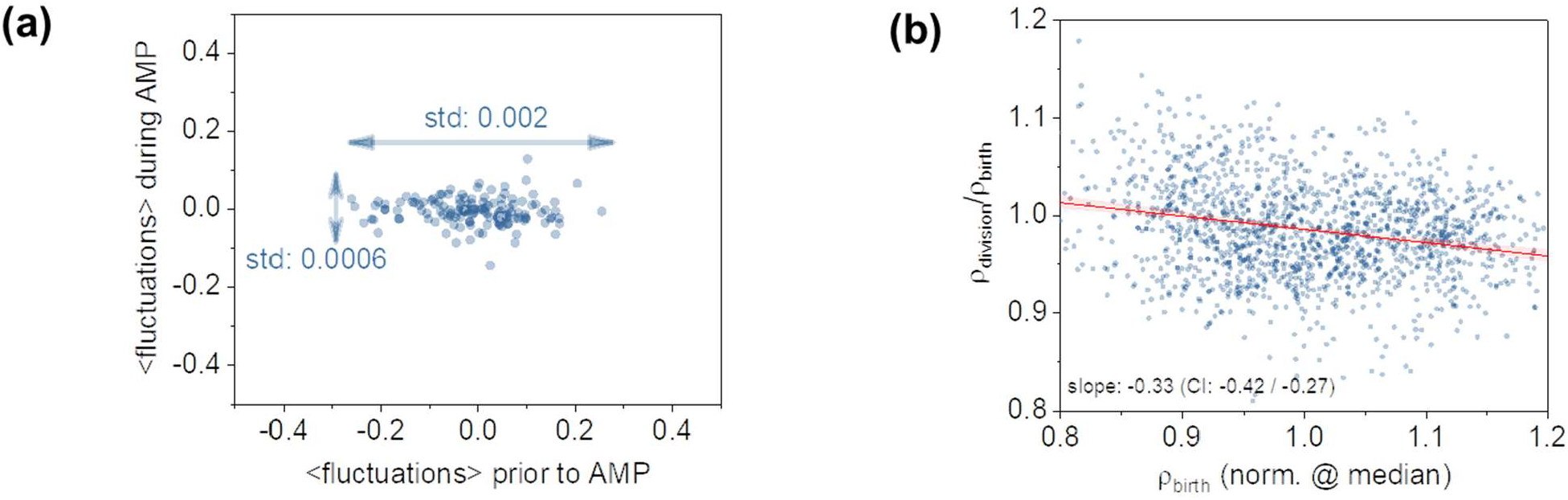
**(a)** Decrease of density fluctuations under the ampicillin (AMP) pressure; legends denote the standard deviation of fluctuations before (*horizontal arrow*) and during the ampicillin treatment (*vertical arrow*); graph represents the cumulative response of three replicates with each replicate presented separately in **Sup. Fig. 8. (b)** Density homeostasis, with cell density prior to division (ρ_division_) being dependent on the cellular dry-density at birth (ρ_birth_); graph plots the combined three replicates, with each replicate presented separately in **Sup. Fig. 9**.

Intrigued by these findings, we explored whether a density homeostasis mechanism may be at play. To this end, we linked the median density fluctuations during growth (dρ/dt) with the resulting differentiation behavior and the ratio of birth and division densities (ρ_division_ / ρ_birth_). We observed that cells exhibiting γ_A_ > γ_M_ reached division with lower dry-density than their density at birth (ρ_division_ < ρ_birth_, **Fig. 2b**); conversely cells exhibiting γ_M_ > γ_A_ concluded their cycle with higher dry-density at division than at birth (ρ_division_ > ρ_birth_, **Fig. 2b**). We reasoned that this observation represents a density homoeostasis mechanism, where density fluctuations maintain cellular density closer to the population average. To investigate this homeostasis mechanism, we considered a simple mathematical model of density fluctuations (dρ/dt) during growth. Based on the notion of exponential size (A) and mass (M) growth, then density dynamics can be expressed as *dρ/dt = (γ*_*M*_ *– γ*_*A*_*)×t* (**Methods**). If γ_M_ and γ_A_ do not depend on density then the above model is not homeostatic (even when γ_M_ = γ_A_), as the slightest noise in γ_M_ or γ_A_ would enforce density fluctuations to grow unboundedly^53, 54^. In contrast, density homeostasis arises by making γ_M_ and γ_A_ density dependent. We experimentally verified this dependence, as evidenced by the monotonic decrease of γ_M_ - γ_A_ with respect to the newborn cell density (**Fig. 3b**). The slope of this function was negative (−0.33 with a [-0.42, -0.27] 95% confidence interval, determined by bootstrapping, **Methods**), reflecting the control of cellular density in the form of negative feedback^55^.

### Fitness Effects

Finally, we explored how density fluctuations and the resulting growth differentiation may confound γ_A_ and γ_M_ as predictors of single-cell fitness, as previously postulated^16, 23^ or expected from population-level Malthusian models^16, 37^. Here, we quantify the fitness of a single-cell through the rate of (asexual) production of progeny, or alternatively the inverse of the life-cycle duration, as also posited by others^56^. We note that this definition pertains to the fitness levels of a single-cell rather than that of the population^57^. Further, this definition coincides with the time required by a cell to complete its cycle as typically reported in size control and homeostasis investigations^21^. Similarly, this definition captures the notion of the instantaneous reproduction capability of single-cells and the rate of gene contribution to the next generation in a constant environment. By experimentally delineating fitness, γ_A_, and γ_M_, we observed that fitness generally increased with γ_M_ and γ_A_ (**Fig. 4a**), with γ_A_ displaying a moderately stronger effect (evidenced by the higher slope in the relationship of γ_A_ – fitness than γ_M_ – fitness, **Sup. Fig. 10**); this effect, however, was not statistically significant due to the overlapping confidence intervals and, importantly, the several low fitness individuals persisting at high rates of size and mass accumulation (**Fig. 4a**).

**Figure 4.**
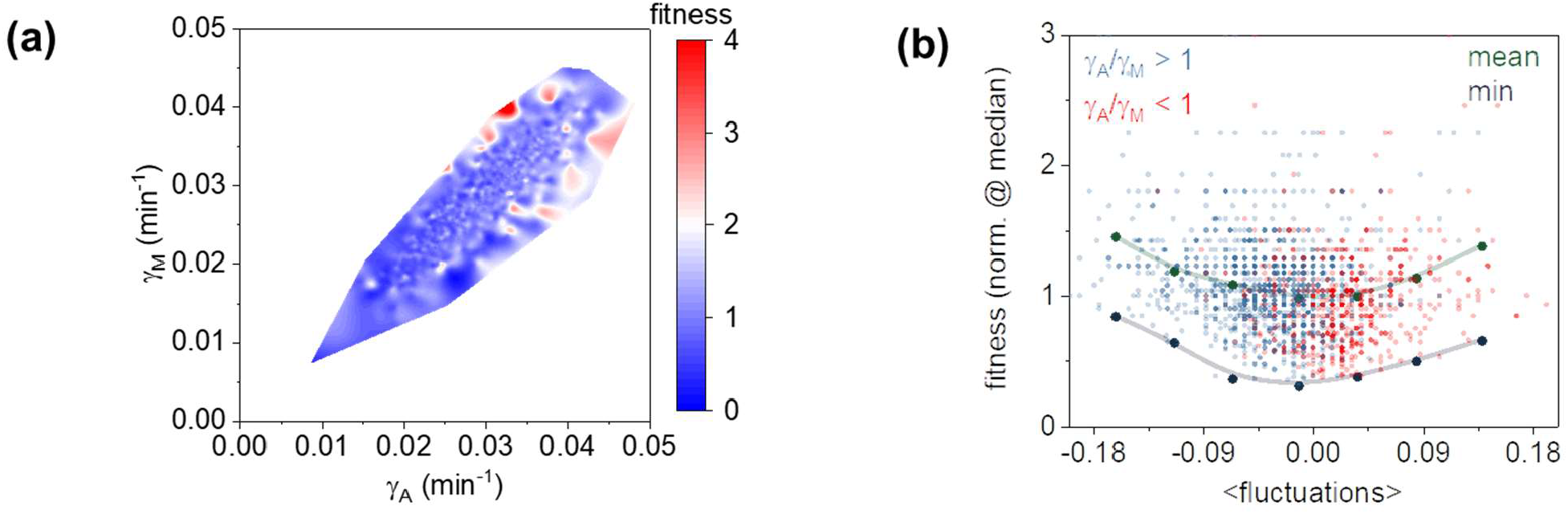
**(a)** 3D representation of growth differentiation (γ_A_ - γ_M_ relationship), with each single-cell observation color-coded by its fitness level; graph represents the cumulative response of three replicates with each replicate presented separately in **Sup. Fig. 13. (b)** Fitness plotted as a function of density fluctuations; blue and red data points represent single-cell observations (color coded by their level of differentiation); green and purple points represent the averaged binned data and minimum fitness levels at different levels of fluctuations; similarly, graph plots the combined three replicates, with each replicate presented separately in **Sup. Fig. 14**.

Unexpectedly, we observed instead that increased levels of differentiation could predict high fitness levels (**Fig. 4b**). In this context, increased density fluctuations (either positive or negative) enabled higher fitness levels even at lower γ_M_ and γ_A_ levels (**Fig. 4b**). Similarly, low fitness individuals were substantially less abundant at increased fluctuations levels (**Fig. 4b**). This observation indicates that increased fitness does not generally require high γ_A_ (as previously postulated^23^) or γ_M_ (as expected from the population-level Malthusian parameter^37^), but can also be expressed by individuals that undergo increased density fluctuations. On the one hand, such individuals may be born with high density that may be accompanied by the gratuitous overexpression of the components required for growth (e.g., ribosomes)^5, 58^. On the other hand, cells may be born with low-density that, mechanistically, may invest more on mass production rather than size accumulation.

## Discussion

In summary, we report that while individual *E. coli* cells accumulate size and mass at exponential rates, their dry-density fluctuates non-monotonically during growth. These density fluctuations not only enable key insight into the innate cell-to-cell density variability in clonal populations (**Fig. 1a**), but also establish that single-cells grow at imbalanced rates of size and mass accumulation. Specifically, we observed that depending on whether density fluctuations are, on average, positive (or negative), clonal subpopulations emerge that optimize mass accumulation with respect to size (and vice-versa).

We linked this form of growth differentiation with the tendency of cells born at a density that is different than the population median to accordingly adjust their density. In this context, cells exhibiting higher rates of mass production reach division at a higher intracellular density state (**Fig. 2c**). Conversely, cells born at a density that is higher than the population’s, exhibit higher rates of size increase, therefore decreasing their density at birth. These phenomena are inhibited upon division inhibition, forming a previously masked density homeostasis mechanism, namely the maintenance of cellular density close to the population average. We validated this mechanism with a simple mathematical model. Further, we demonstrate that both single-cell size and mass of *E. coli* are consistent with the adder phenotype^19, 21, 22^ (**Sup. Fig. 11** and **12**), It is worth noting, however, that the adder model does not take the observed density fluctuations into consideration, suggesting the presence of an additional control layer that is not represented by the adder phenotype.

Importantly, we report a previously masked link between growth and fitness at the single-cell level that does not conform with previous single-cell postulates^23^ and population-level (e.g., Malthusian) models^37^. Specifically, we found that individual cells that exhibit high levels of fitness can be better identified by their density-fluctuations amplitude than the respective rates of size or mass accumulation (**Fig. 4a**). In this context, we observed that high (low) fitness individuals were substantially more (less) abundant at increased fluctuations levels (**Fig. 4b**). Mechanistically, this can be thought of as gratuitous overexpression of growth-required molecular components at birth, specifically for individuals undergoing increased negative density fluctuations; conversely, overall positive density fluctuations of increased levels may enable cells to invest more in mass production than size accumulation, thereby reducing the time required to reach division.

Finally, we detail our experimental approach that enabled the dynamic tracking of cellular density with high precision and throughput rates, which is challenging, if not impossible, with conventional microfluidic systems. For enhanced precision, we fused quantitative-mass imaging with 1D microarrays that not only deterministically eliminated cell crowding, but also became invisible upon contact with water. This approach minimized light scattering at the cell-to-cell and cell-to-microfluidics interfaces, thus, preserving key optical information during microcolony expansion (**Fig. 1b**). We also employed advanced microfabrication to improve the underlying throughput rates by more than one order of magnitude in comparison to existing assays. These novel methodological approaches can be directly translated to alternative species to further explore the role of cellular dry-density and related effects in growth, division, and fitness of single-cells. Given that all inoculants and infections start from a single, or very few growing cells, we anticipate that the paradigm of density fluctuations will improve synthetic, systems, and evolutionary biology investigations that take the segregated notion of non-genetic variability into consideration^59^.

## Materials and Methods

### Strains, Growth Conditions, and MIC levels

#### Strains

Two strains were used in the reported investigations, namely: *Escherichia coli* DH5α (WT) and the ampicillin-resistant *E. coli* E212K mutant, also derived from DH5α. This derivative was chosen to include one more strain in our density fluctuations observations (i.e., in the absence of antibiotics pressure), and specifically carries the g628a mutation (using Ambler numbering^60^) on the TEM-1 gene on pBR322.

#### Growth conditions

WT and resistant strains were grown in batch using a bath incubator (C76, New Brunswick Scientific) at 37oC and 180 rpm. As a growth medium, we employed the Mueller Hinton broth (Difco 275730, BD). The strains were first passed from agar plates (stored at 4oC) to 5 ml medium (round bottom polystyrene tubes, VWR) until early stationary phase, then diluted in 20 ml fresh medium (125 ml glass flasks, Corning) at a 0.01 optical density (OD_600_, λ = 600 nm, V-1200 spectrometer, VWR), and incubated for 12h (37°C, 180 rpm). Overnight cultures were diluted to an OD_600_ of 0.01 (in 20 ml fresh medium), re-grown to mid-exponential phase (∼3 h), and sampled to perform all reported single-cell experiments. We employed the same procedure to determine the MIC levels of the E212K mutant, as detailed below. All single-cell experiments were performed in triplicates by repeating the abovementioned procedure on different days.

#### Minimum inhibitory concentration (MIC)

We measured the MIC levels of E212K using the microdilution method^61^. Following growth (see above), cultures were diluted to 10^6^ cfu/ml (∼0.002 OD_600_) to a volume of 1 ml and transferred to 1 ml of ampicillin (VWR0339, VWR) solution in Mueller Hinton at concentrations ranging from 8,196 to 0.0156 μg/ml at a 1.4× step size^62^, including a 0 μg/ml control. The suspensions were incubated (37°C, 180 rpm) for 20 h to determine the MIC level, namely the lowest ampicillin concentration yielding zero OD^61^, at 176 μg/ml (**Sup. Fig. 15**). We applied this value in all subsequent single-cell experiments. The measurement was repeated (3 times mid-exponential phase cultures and 4 times using stationary phase cultures) yielding the same result.

### Single-Cell Assays

Single *E. coli* cells were laterally confined using 1D microarrays and vertically confined via a top-integrated membrane (**Sup. Fig. 1**), enabling size, density, and mass tracking of single-cells for up to 6-7 generations. The 1D microarrays were fabricated by electron beam lithography in SU8, subsequently transferred to PDMS, and then to a UV curable polymer that was index matched to water (Bio-133, My Polymers). Nutrients were provided by a doped membrane or a microfluidic channel. The latter was applied in the reported antibiotic experiments to yield dynamic switching between medium and ampicillin conditions. The assay microfabrication, assembly, and nutrient supply are further detailed in the Supplementary Information.

### Imaging

#### Cell growth

Quantitative-mass imaging was performed using a spatial light interference microscopy system (Cell Vista Pro, Phi Optics) integrated with an inverted microscope (DMi8, Leica) equipped with an automated stage. In our SLIM system, the quantitative-phase images are formed by projecting the back focal of the imaging, phase-contrast objective onto a liquid crystal spatial light modulator (SLM). The SLM exhibits the appropriate ‘ring-shaped’ phase masks that shift the optical-phase of the light wavefront scattered by the sample relative to the un-scattered light, as detailed in the related original report^38^. In this way, images representing the relative phase delay of E. coli cells (scattered wavefront) with respect to the background (un-scattered wavefront) are formed. 3D z-stack images (0.5 3 μm step size) were acquired using a 63× (NA 0.7, PH2) or 3D z-stack images (0.5 μm step size) using a 40× objective (NA 0.6, PH2) and a 3.65 μm pixel CCD camera (GS3-U3-28S4M, Point Grey Research). To correct for halo effects, and in addition to arranging the microcolonies in 1D as detailed earlier and presented in **Fig. 1c** and **Sup. Fig. 2**, we also processed the quantitative-phase images to remove any residual halo using the procedure described elsewhere^38^. This step increased the background uniformity and enabled moderately better definition of the cell contour, which is key for cell segmentation^63^. A comparison of a single-cell phase image with and without additional halo correction is displayed in **Sup. Fig. 16**. Various locations were imaged every ∼3 minutes (every ∼5 mins for the ampicillin experiments) using automated routines (Metamorph, Molecular Devices). All single-cell growth experiments were performed in triplicates yielding a total of n = 1520 observations of dividing cells, with each replicate consisting of 442 (*rep. 1*), 573 (*rep. 2*), and 505 observations (*rep. 3*). These observations (and related analyses) exclude the very first mother cell. Specific to the dry-density comparison between mothers and daughters (**Fig. 2d** and **Sup. Fig. 7**), the ratios were calculated by considering dividing mothers, and both dividing and non-dividing daughters, thus, enabling the consideration of all diving mothers, yielding 412 (*rep. 1*), 582 (*rep. 2*), and 452 (*rep. 3*) observations. Finally, in the ampicillin (AMP) experiments we did not consider cells with fewer than 4 observations (both prior to and during AMP pressure), as well as non-growing cells (e.g., persisters), yielding 60 (*rep. 1*), 45 (*rep. 2*), and 35 (*rep. 3*) observations.

#### Throughput analysis

Here, throughput denotes the number of observations per field of view, or alternatively the maximum possible number of (4 generations) microcolonies in a single image. We compared the 1D and 2D assay throughputs at 20× (NA 0.4, PH1) and 40× (NA 0.6, PH2) magnification, respectively using quantitative-phase imaging and a 6.5 μm pixel size sCMOS camera (ORCA-Flash 4, Hamamatsu). For 1D, we imaged ∼1,000 fixed DH5α cells (overnight fixation in 2.5% glutaraldehyde at 4oC, followed by 3× washing in medium, and diluted at a varying density from 0.01 to 0.175), which we introduced in the 1D microarrays. We followed a similar approach in 2D, albeit using live cells that we allowed to grow to microcolonies containing ∼20 cells. This was performed in triplicates, with each replicate containing 40 microcolony observations.

### Image Analysis

#### Cell growth

All quantitative-mass images were processed using ImageJ and Fiji (National Institutes of Health), Metamorph, and MATLAB (Mathworks), as follows: **(1)** choice of the best focus plane (P_i_) and maximum projection of P_i_, P_i-1_, and P_i+1_; **(2)** filtering by median and gaussian blur (ImageJ), 2D deconvolution (*No Neighbors*, Metamorph), and 1D Fast Fourier Transform (FFT, ImageJ); and **(3)** thresholding via the Maximum Entropy algorithm. All resulting binary images were subsequently subjected to watershed, visual inspection, and – if necessary – manual curation. Following processing, the binary and original quantitative-phase images were assembled into two separate time-lapse stacks, divided into microcolonies, and analyzed with lineage mapper (Fiji) to extract lineage trees and track single-cells from birth to division.

#### Assessment of Image Processing

We selected the abovementioned image processing pipeline for its robustness and reduced computational requirements, as we have previously demonstrated for bacteria and yeast^31, 63^. Further, while density fluctuations have been reported by others^34^ and we also observe them in high temporal resolution readings (with smoother traces, **Sup. Fig. 17**), we performed additional steps to ensure that our observations are not due to cell segmentation or plane selection errors. Specific to cell segmentation, we ensured validity by performing the following two analyses. *First*, we compared our abovementioned with the Otsu and Moments thresholding algorithms. *Second*, we compared our approach to a 1D segmentation approach that is independent of conventional thresholding algorithms and, thus, possible errors in area segmentation. In this context, we determined the beginning and end of a 1D cell-contour (of a constant 6-pixel width) at 20% above the noise floor. All comparisons (**Sup. Fig. 17**) yielded moderate differences in the single-cell density dynamics, which upon normalization (at t = 0 or the time of birth) exhibited very high agreement in single-cell density dynamics characterized by greater than 99% Pearson correlation coefficients (p < 0.001). This finding suggests that the observed density fluctuations represent a physiological response, largely independent of potential errors during cell segmentation.

Further, we ensured that we selected the proper plane of focus by inspecting the quantitative-phase images of all cells at all collected z-planes, as well as the z-dependence of their phase signal. To this end, we employed a custom MATLAB code that simultaneously displayed cell images of all planes and selected the plane (P_i_) that exhibited the sharpest image. Following maximum projection between P_i+1_ and P_i-1_, we visually inspected all images to appropriate plane selection. Furthermore, we estimated the induced density uncertainty after intentionally selecting a wrong plane of focus. To this end, we intentionally selected ± 1 plane away from focus and computed the resulting single-cell density error (standard error). In this way, we computed a 0.78% uncertainty in the density determination of a single-cell due to an experimental error in plane selection, which is lower than the observed density fluctuations, as displayed in **Sup. Fig. 17**.

#### Throughput analysis

ImageJ was used to quantify 1D and 2D throughput using the previously detailed procedures. To quantify 1D throughput, we used the resulting statistics of 1,000 cells to determine the average cell length and the distance of each cell to its nearest neighbor. To quantify 2D throughput, we analyzed images of microcolonies containing up to 16 cells to determine the largest microcolony dimension using the Feret’s diameter (ImageJ). In this context, we did not approximate the microcolony as a circle, given that 2D confined *E. coli* microcolonies are known to form dynamic nematic patterns of variable asymmetries and stochastic orientation^64^. Our observations denoted the same effect (**Sup. Fig. 2b**). To compare information loss in 1D and 2D, we analyzed images of 1D and 2D microcolonies containing 20 single-cells. Subsequently, we averaged the dry-density of all cells in the microcolony, as reported in **Fig. 1c**.

### Data analysis

#### Cell growth

The following metrics were extracted from each image: area, density, and mass per time-point per cell. Growth rates were computed using these functions of *A*(*t*) = 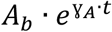 and 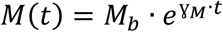 in MATLAB throughout the cell cycle. Density (ρ) fluctuations were determined as the median of dρ/dt, where dρ represents the ρ(t_i+1_) - ρ(t_i_) difference in the dt = t_i+1_ - t_i_ window. To quantify the density of single-cells from the measured optical phase delay, we used the following expression^30, 31, 39^:

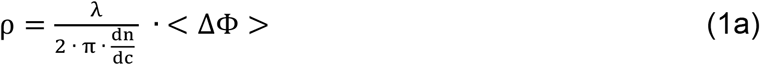

with 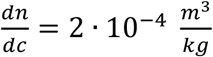 representing the protein-specific refractive index increment ^26^, λ the wavelength of illumination (centered at 550 nm), and <ΔΦ> the average experimentally determined phase delay difference between the cell cytosol and the extracellular medium, integrated across the cytosolic area A. To compute the cell mass, we multiplied cell density with its area. We note that eq. 1 computes the area-based density of single-cell without essentially requiring prior knowledge of the cell area. This approach have been previously demonstrated in^30^ and^39^.

#### Throughput analysis

1D throughput was quantified via a nearest neighbor analysis (MATLAB, *knnsearch, euclidean*). Specifically, we set the minimum distance to the nearest neighbor equal to the average cell length multiplied by 16 (i.e., the expected number of progeny in a microcolony for the duration of our experiments). In this context, any individual cell exhibiting horizonal distances (i.e., in an axis parallel to growth) from the nearest neighbor that were greater than this threshold was taken into consideration. We followed a similar procedure for quantifying 2D throughput. Here, after determining the largest microcolony size through the Feret’s diameter, we performed a 2D nearest neighbor analysis (MATLAB, *knnsearch, euclidean*) using the Feret’s diameter as a threshold.

### Models

#### Density Homeostasis

Let mass *M* of a single cell grow exponentially with rate γ_M_ during the cell cycle:

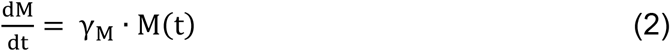

Similarly, area *A* of a single cell grows exponentially with rate γ_A_:

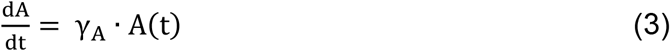

Then, density (defined as the mass over area ratio) evolves as:

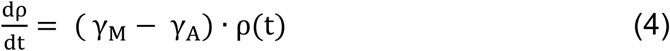

If γ_M_ and γ_A_ are density independent then the above model is not homeostatic even when γ_M_ = γ_A_, as the slightest noise in these rates makes density fluctuations grow unboundedly over time ^53, 54^. In contrast, density homeostasis arises by making γ_M_ and γ_A_ density dependent. Let

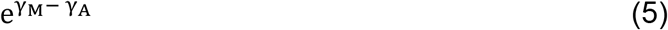

be a monotonically decreasing function *f* of newborn cell density ρ_i_:

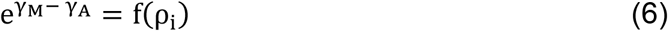

As such, a newborn with a low density will invest more in mass growth *vs*. area growth. Substituting (6) in (4), the density at the end of cell cycle is:

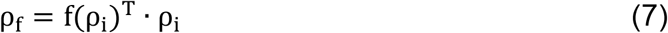

where *T* is the length of the cell-cycle. With this, one can write the following iterative model for the newborn densities *ρ*_*i*_ in the *n*^*th*^ generation:

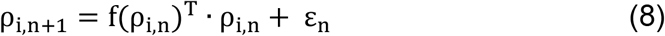

Where ε_n_ is the noise induced at division from random partitioning of area and mass. The above model has a unique fixed point given by the solution to the equation:

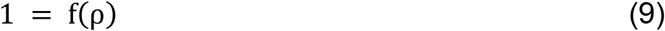

which will be a stable homeostatic set point for ρ in the presence of noise as long as the specialization function *f* is a decreasing function of density. Eq. 6 can be rewritten as:

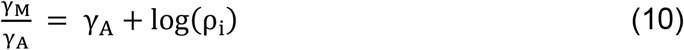

Equation 10 implies that the γ_M_/γ_A_ ratio within a cell cycle should also be decreasing function of the newborn cell density, as was experimentally observed and displayed in **Fig. 3b**.

Using some of the ideas put forward by Oldewurtel et al.^34^, we further explored potential mechanisms that could underly density homeostasis. One such mechanism includes:

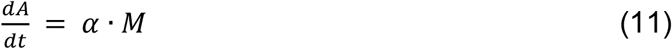

This equation denotes that part of the proteome is dedicated to the increase of cell size. This notion is also congruent with density homeostasis. In this context, Eq. 1 leads to the following expression for the temporal evolution of density:

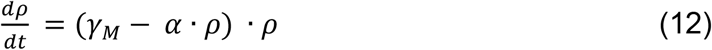

In this case, the density at steady state is given by:

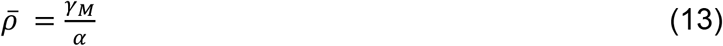

Solving the differential equation (2), the ratio of densities at the start and end of cell cycle is

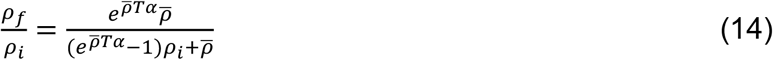

where *T* represents the duration of the cell-cycle (or cell fitness). This ratio is a decreasing function of ρ_i_, as noted in **Fig. 3b**, which suggests density homeostasis.

Given that both mass and area increase exponentially as per

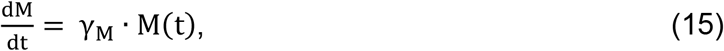

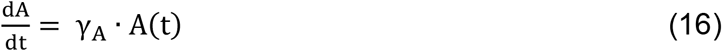

Then the model described by equation (11) corresponds to γ_A_ being an increasing function of density. Indeed, this is the behavior we observe in our experimental data, as evidenced in **Sup. Fig. 6**.

An alternative model of density of homeostasis, also presented by *Oldewurtel et al*,^34^ is:

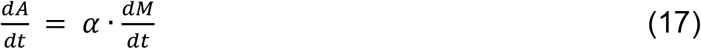

This expression corresponds to the ratio γ_A_/γ_M_ being proportional to density at birth. While we do see a modest increase in γ_A_/γ_M_ as a function of density at birth, we observe a stronger increase with γ_A_ as function of density (**Sup. Fig. 6**). These differences suggest that the model presented in Eq. 11 is likely the dominant driver of density homeostasis.

#### Adder, sizer, and timer

To compare cell size at division (A_d_) with the adder, sizer, and timer models from the size at birth (A_b_), we employed the following expression ^21^:

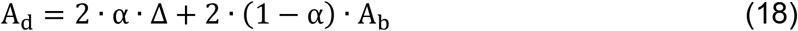

We adapted eq. (11) to similarly represent cell biomass at division (M_d_) as:

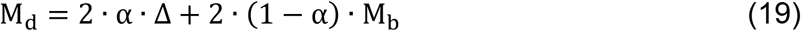

In both equations, α differs as: α = ½ for adder, α = 1 for sizer, and α = 0 for timer. Δ is the median area (A_b_) and mass (M_b_) at birth. The results of these three models are plotted in **Sup. Fig. 11** (for size) and **Sup. Fig. 12** (for mass) and compared to the experimental raw data (scatter plot), a linear regression (based on the experimental data), and the binned experimental data.

### Statistics

The robust coefficient of variation was derived in MATLAB using the *mad(X,1)* function to first calculate median absolute deviations that we divide it with the population’s median. ANOVA tests were performed in MATLAB using the *anova1* function. The 95% confidence intervals of the linear regressions were computed in MATLAB by bootstrapping using the *bootci* function (n = 1,000 samples). Binning was performed in MATLAB using the *Sturges* method and the *histcounts* function. Mann-Whitney, Kolmogorov-Smirnov, two sample t tests, and all plotted linear regressions were performed in Origin Pro.

## Acknowledgments

AEV and AMD acknowledge support from the U.S. Department of Energy, Office of Science, Office of Biological and Environmental Research (DE-SC0019249). SN, DW, and AEV acknowledge support from the National Science Foundation (OIA-1736253). AS acknowledges support from the National Institutes of Health (5R01GM124446). Discussions with Larry Forney are also gratefully acknowledged.

## Author Contributions

SN performed the imaging experiments, image and data analysis, and contributed to the manuscript preparation; AS developed the analytical homeostasis model; SDD and AMD contributed to the microfabrication of the assays; DW contributed the strains; AEV conceived and overviewed the research, and wrote the manuscript.

