## Supplementary Figures for "Density Fluctuations Yield Distinct Growth and Fitness Effects in Single Bacteria"

**Supplementary Figure 1**

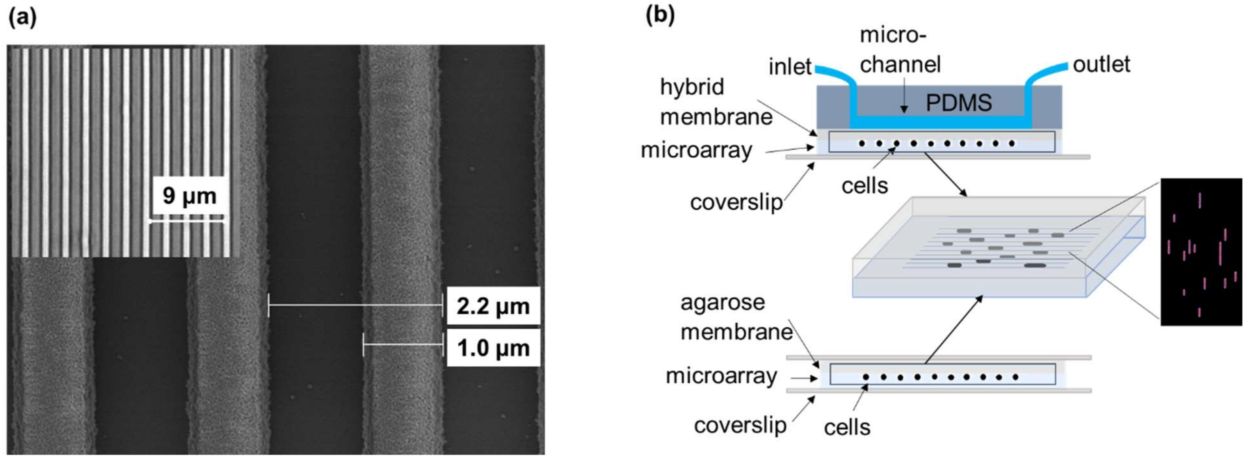


**(a)** Scanning Electron Microscopy (SEM) of a region of the 1D microarray in SU8 on a Si wafer (~1.2 µm linewidth where cells reside in the 1 µm spacing); *inset* displays a bright field image of the same microarrays transferred onto the BIO133 polymer. **(b)** Schematic representation of the 1D growth microarrays; *top*: cells positioned in the 1D microarrays and vertically confined via a hybrid membrane and a microfluidic system that delivers medium to the cells; *bottom*: cells positioned in a 1D microarray and confined vertically by a top-integrated agarose membrane that is doped with nutrients; *middle*: 3D representation of the cell-growth region. In all cases, imaging was performed through the bottom coverslip.

**Supplementary Figure 2**

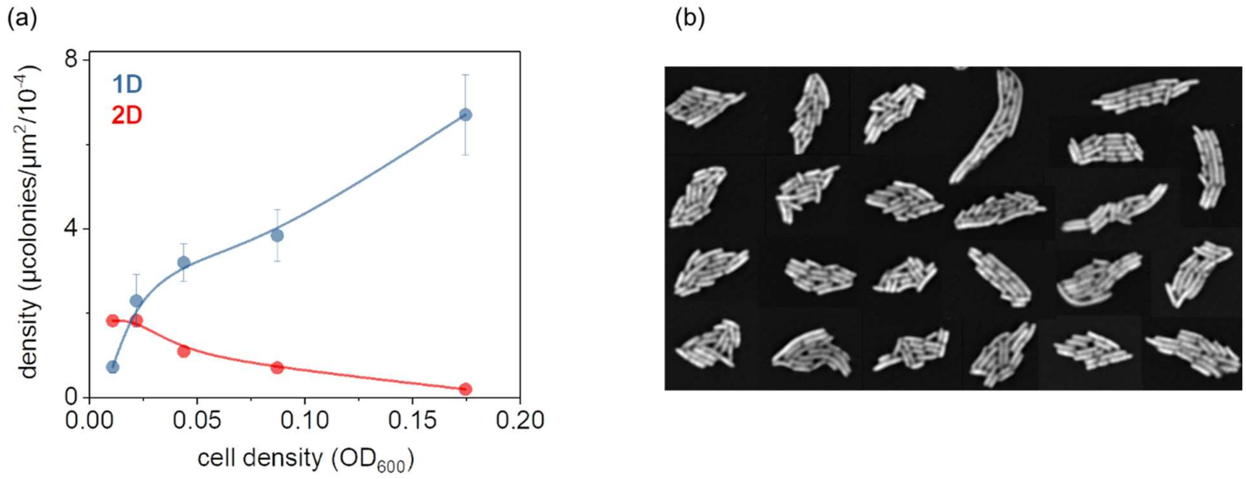


**(a)** Throughput (i.e., number of observations per unit area) of the 1D (at 1 µm spacing) and 2D growth assays as a function of the optical density of the deposited samples (color-coded); data points represent the average and error bars the standard deviation of 3 replicates for growth of up to 16 cells; the microcolony density per unit area is significantly higher for ODs equal or greater than 0.04 (two-sample t-test under the Welch Correction for ODs 0.044 ($p = 0.009$, $t = 4.78$, DF = 4), 0.088 ($p = 0.007$, $t = 5.03$, DF = 4), and 0.175 ($p = 0.002$, $t = 6.84$, DF = 4). **(b)** A representative montage of several 2D microcolonies illustrating shape asymmetry and underlying stochasticity.

**Supplementary Figure 3**

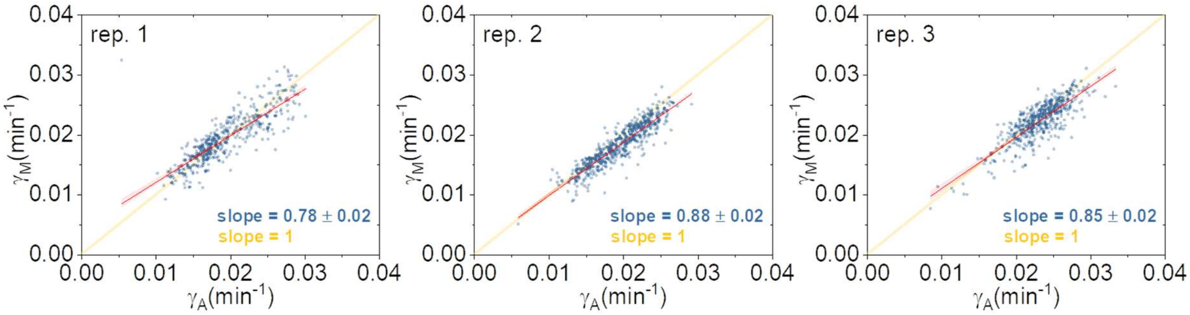


Growth differentiation with some cells optimizing area accumulation rates ($\gamma_A$) and some biomass accumulation rates ($\gamma_M$); each graph corresponds one experimental replicate performed on different days; red line represents the linear fit (shaded areas are the 95% confidence intervals); yellow line represents a hypothetical line of slope 1.

### Supplementary Figure 4

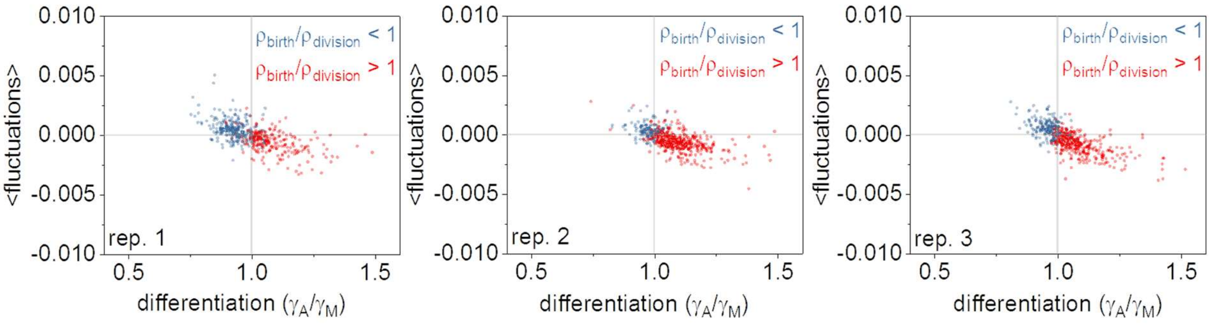


Median density fluctuations (<dρ/dt>) during growth as a function of growth differentiation ($\gamma_A/\gamma_M$) for three replicates; corresponds to overall increasing (blue) or decreasing (red) cellular dry-density during growth.

**Supplementary Figure 5**

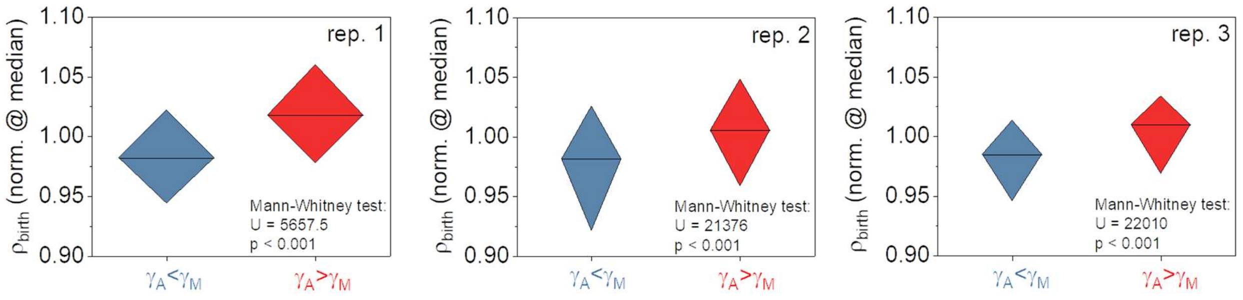


Cells born with lower density than the population median maximize biomass accumulation rates (differentiation or $\gamma_A/\gamma_M < 1$, blue), while cells born with higher than the population median density maximize area accumulation rate (differentiation or $\gamma_A/\gamma_M > 1$, red). Boxcharts represent the 25%-75% of the cumulative response of three replicates; legends summarize the result of the Mann-Whitney test, with additional tests displayed in **Sup. Table 1**).

**Supplementary Figure 6**

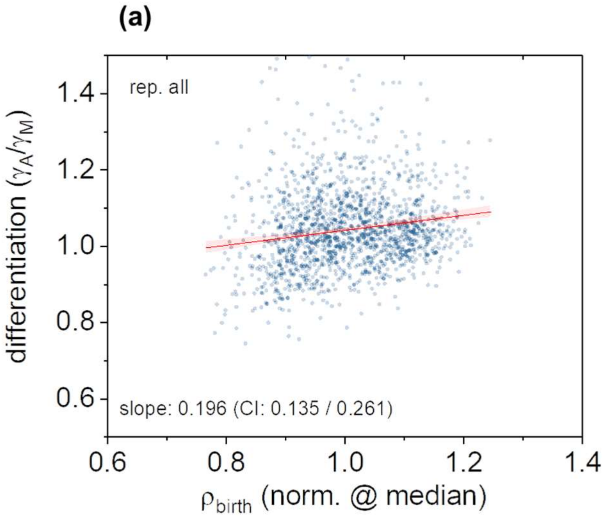


**(a)** Growth differentiation ($\gamma_A/\gamma_M$) as a function of cell density at birth ($\rho_i$). **(b)** Growth rate by size ($\gamma_A$) as a function of cell density at birth ($\rho_i$). In both graphs, all three replicates are combined with the red line representing a linear fit, while legend denotes the linear slope and the corresponding 95% confidence intervals (CI) determined by bootstrapping.

**Supplementary Figure 7**

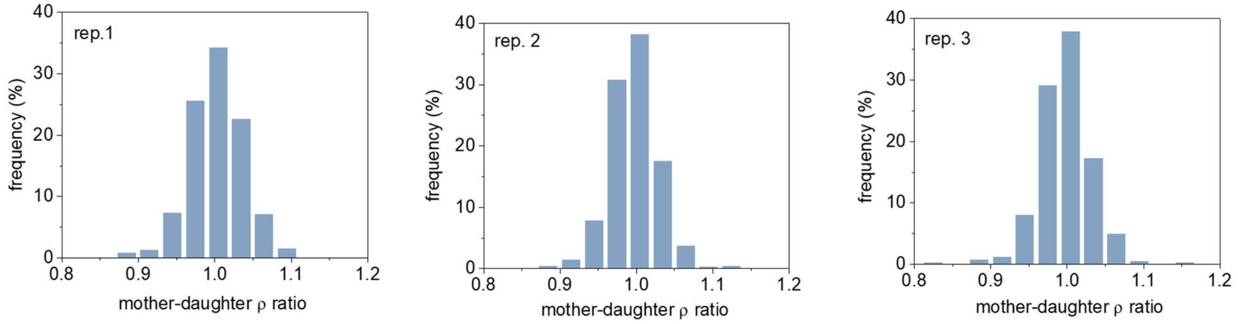


Density similarity between mother cells at division and daughter cells at birth. The graph represents the probability distributions for each experimental replicate separately. Each replicate exhibited comparable robust coefficients of variation (*rep. 1*: 3.4%, *rep. 2*: 3%, *rep. 3*: 2.9%), also comparable with respective value in the cumulative distribution (3.1%).

**Supplementary Figure 8**

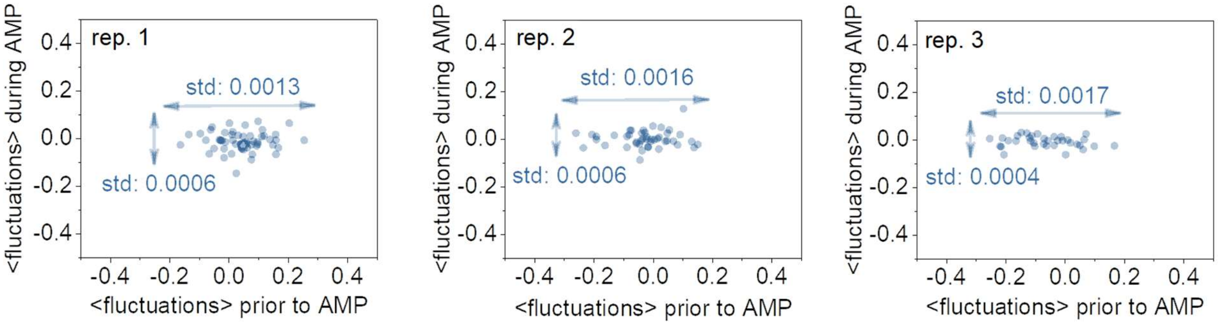


Density fluctuations disappears under the ampicillin treatment; graphs represent three separate experimental replicates; *legends* note the standard deviation of fluctuations before (*horizontal arrow*) and during the ampicillin treatment (*vertical arrow*);

**Supplementary Figure 9**

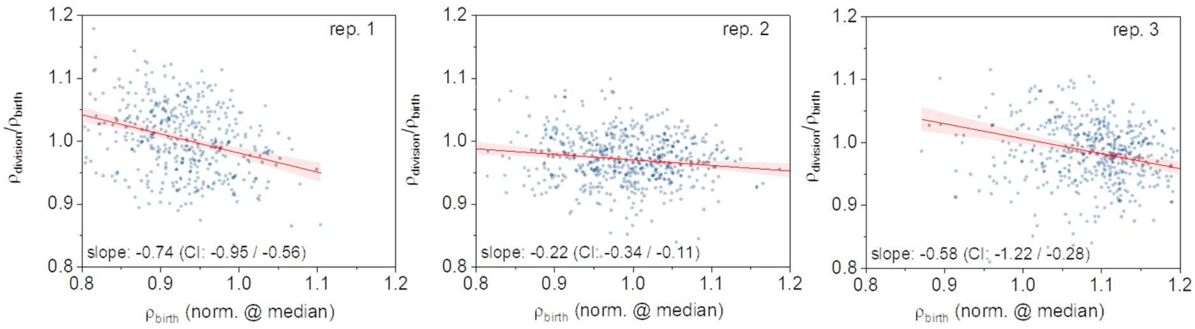


Dry-density homeostasis during the cell cycle, with the final cell density ($\rho_{division}$) being dependent on the cellular dry-density at birth ($\rho_{birth}$); each graph plots the response of each experimental replicate separately.

**Supplementary Figure 10**

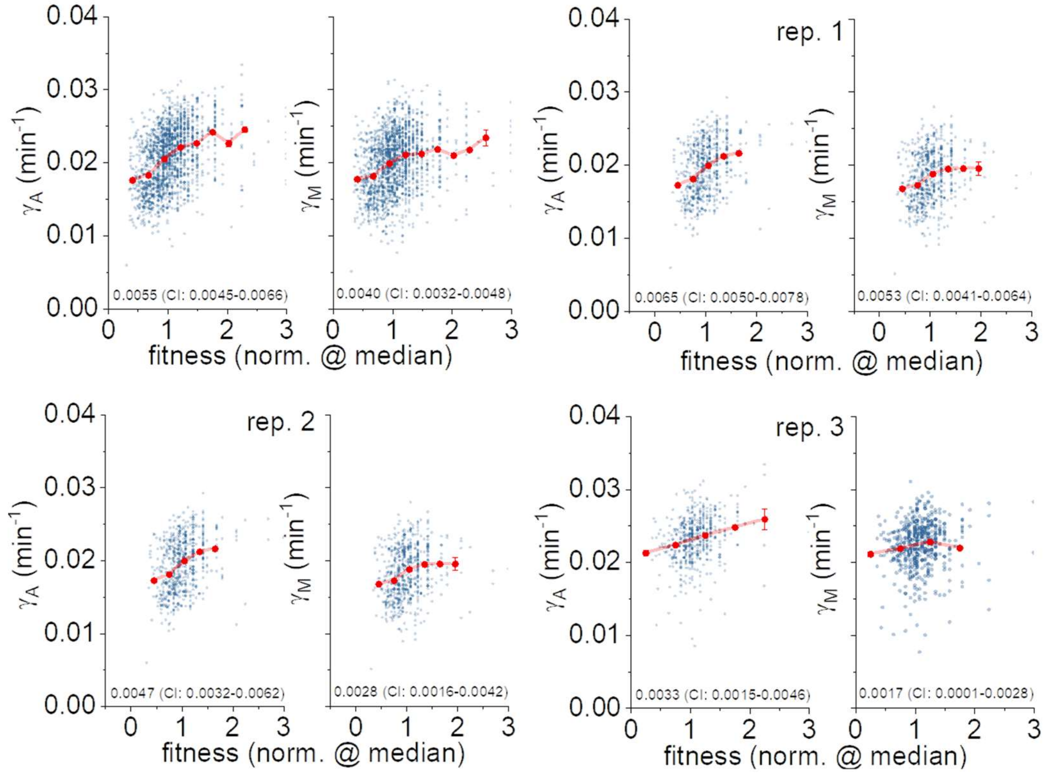


Size ($\gamma_A$) and mass ($\gamma_M$) accumulation rates as a function of fitness for all replicates combined and each individual triplicate; *blue dots* represent the experimental data; *red line* represents the binned data; *legend* displays the slope and the 95% confidence intervals (CI) of the linear fit (by bootstrapping).

**Supplementary Figure 11**

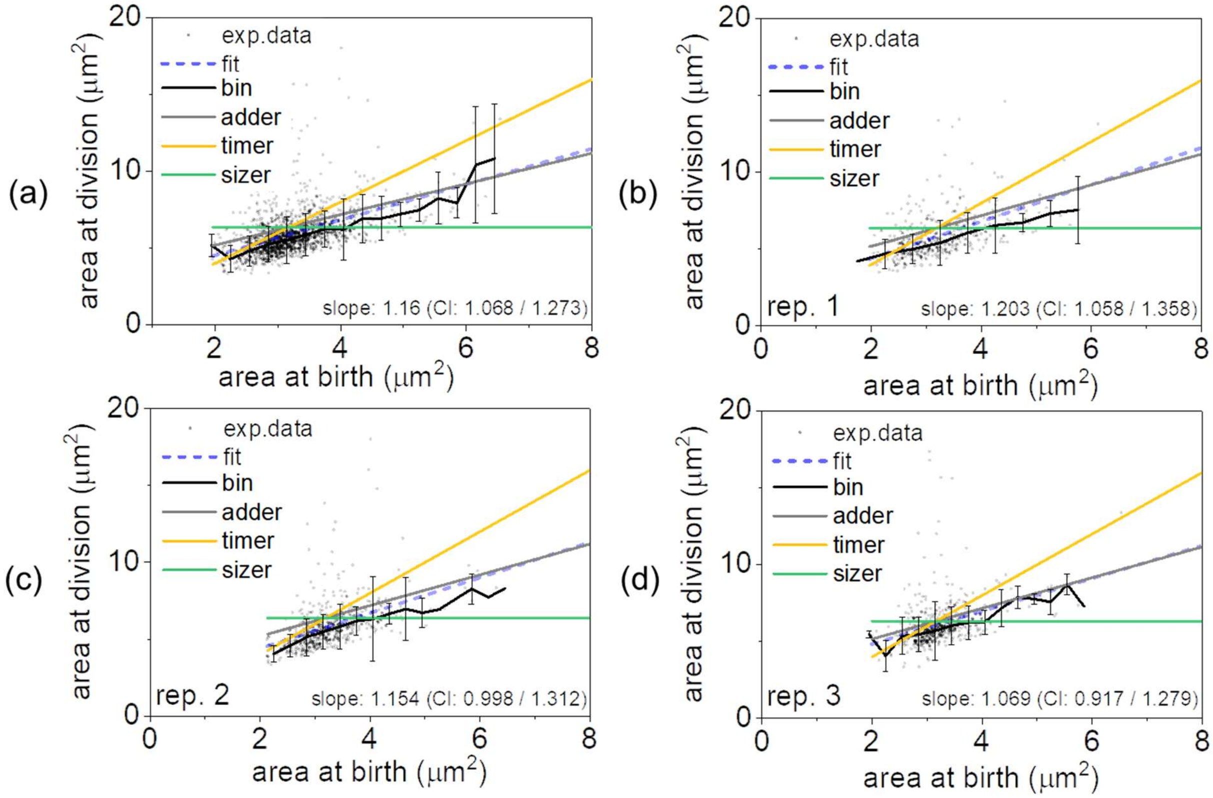


Comparison of the empirical observations of cell size regulation with the adder (*grey*), timer (*yellow*), and sizer (*green*) models. Scatter plots indicate the experimental data; *blue* dotted line represents the linear regression fit (*legend* denotes the slope and the corresponding 95% confidence intervals – CI – determined by bootstrapping); *black* line represents the binning result with the error-bars noting the standard deviation. Clearly, the adder model appears to best fit to the experiment for both the cumulative response of the combined replicates **(a)** and the individual replicates themselves **(b-d)**.

**Supplementary Figure 12**

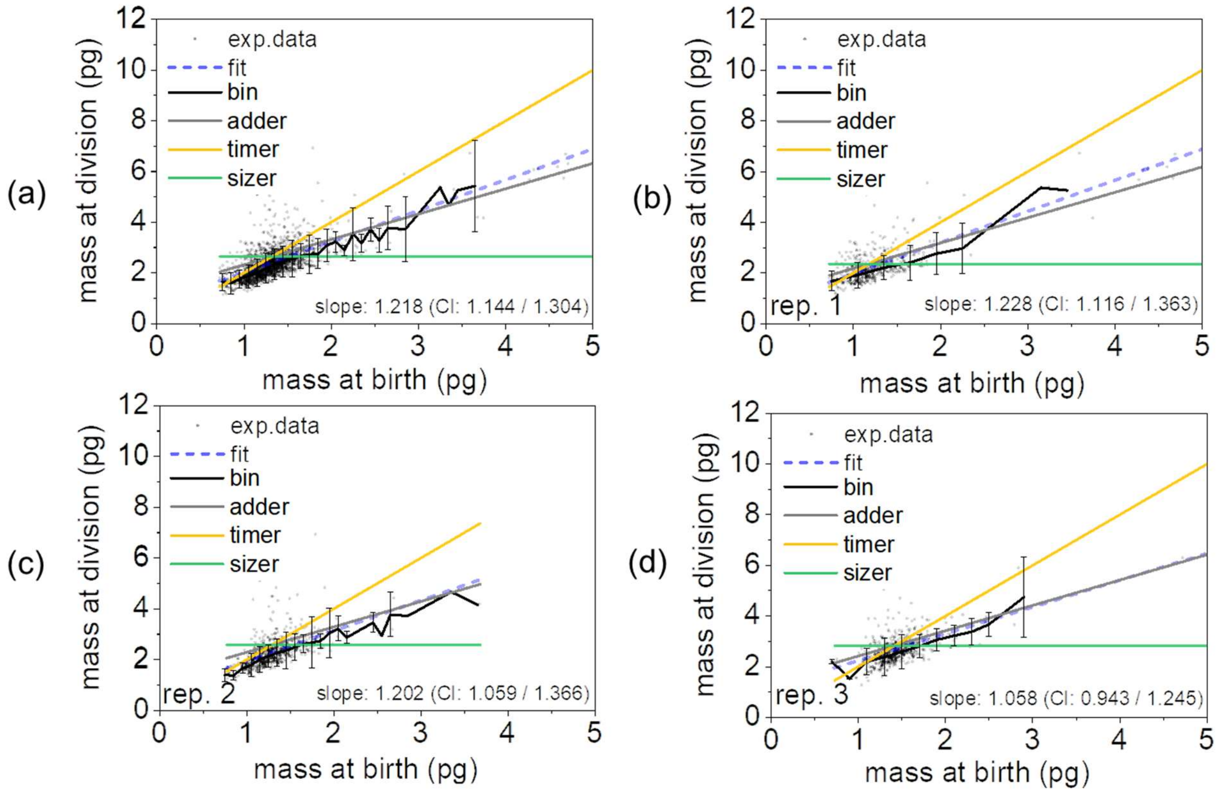


Comparison of the empirical observations of cell mass regulation with the adder (*grey*), timer (*yellow*), and sizer (*green*) models. Scatter plots indicate the experimental data; *blue* dotted line represents the linear regression fit (*legend* denotes the slope and the corresponding 95% confidence intervals – CI – determined by bootstrapping); *black* line represents the binning result with the error-bars noting the standard deviation. Clearly, the adder model appears to best fit to the experiment for both the cumulative response of the combined replicates **(a)** and the individual replicates themselves **(b-d)**.

**Supplementary Figure 13**

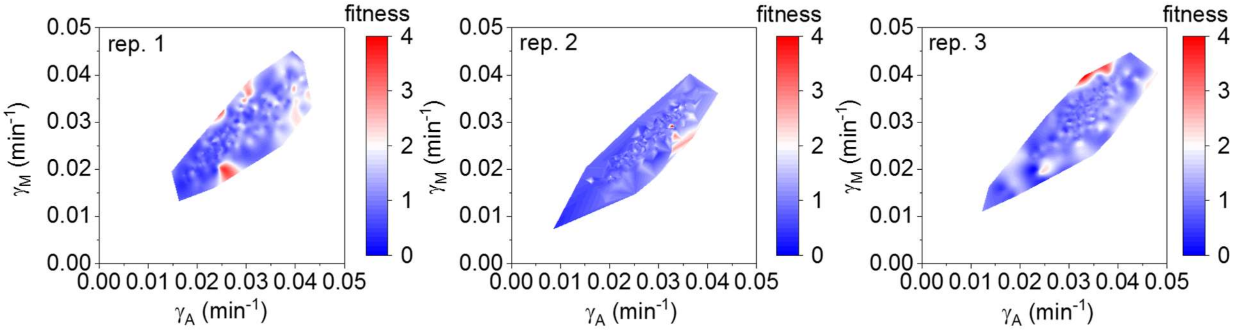


Growth differentiation ($\gamma_A$ - $\gamma_M$ relationship), with each single-cell observation color-coded by its fitness level; each graph displays the response of each experimental replicate separately.

**Supplementary Figure 14**

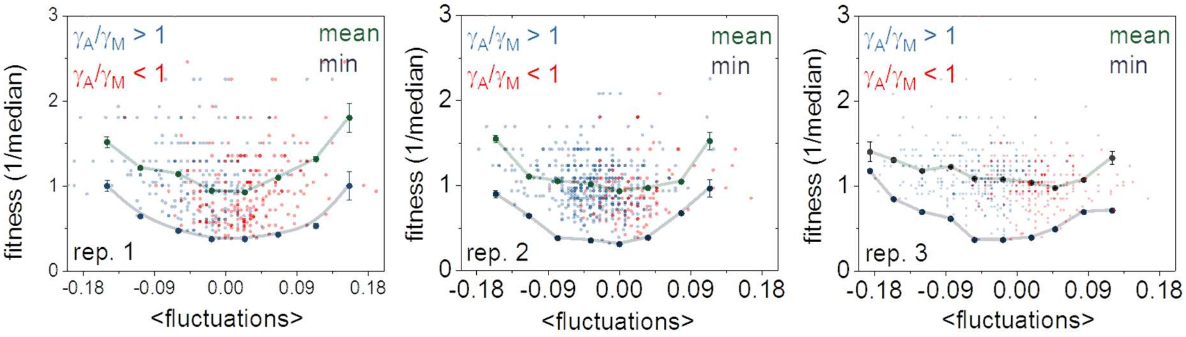


Fitness (normalized to the population median) plotted as a function of density fluctuations; *blue* and *red* data points represent single-cell observations (color coded by their differentiation strategy, $\gamma_A/\gamma_M$); *green* and *purple* points represent the averaged binned data and minimum fitness levels at different levels of fluctuations; each graph plots the response of each experimental replicate separately.

**Supplementary Figure 15**

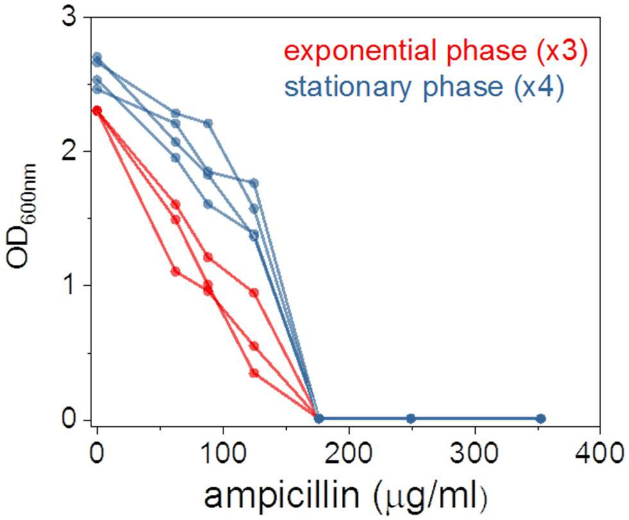


Minimum inhibitory concentration (MIC) determination for the ampicillin resistant E212K mutant using 3 replicates from mid-exponential phase and 4 replicates from stationary phase. No growth was observed at 176 µg/ml ampicillin concentration in all experiments.

**Supplementary Figure 16**

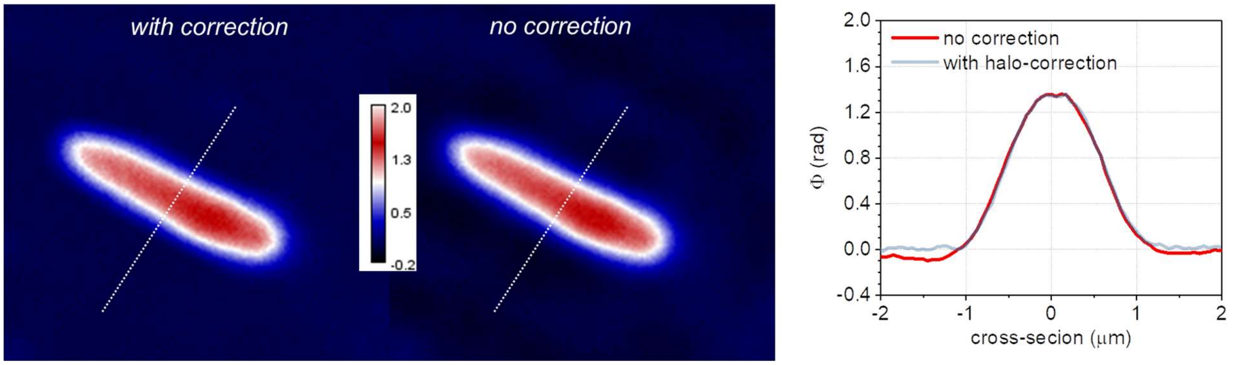


Eliciting the effect of computational halo correction on the phase profile of a single *E. coli* cell using the approach detailed in (42). On the left, the phase images with and without halo correction of the same cell are displayed, while on the right the phase profile along the dotted line is plotted for each image. Halo correction improves the background uniformity and enables better definition of the cell contour, which is critical for cell segmentation.

### Supplementary Figure 17

(a)

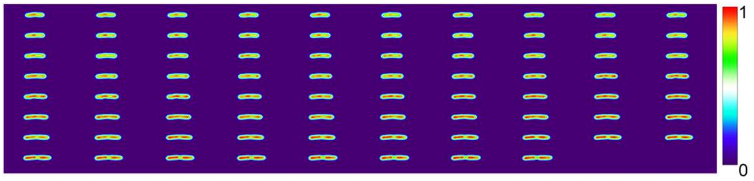


(b)

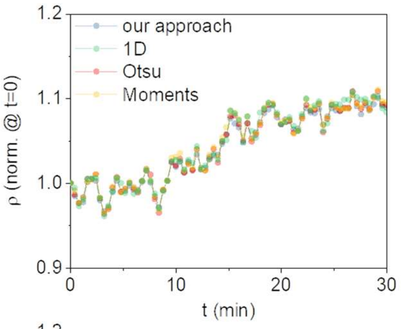


(c)

| Pearson's correlations | result |
|---|---|
| our approach - 1D line | ρ = 0.993, $p = 5.82 \times 10^{-71}$ |
| our approach - Otsu | ρ = 0.996, $p = 1.69 \times 10^{-80}$ |
| our approach - Moments | ρ = 0.995, $p = 2.49 \times 10^{-78}$ |

(d)

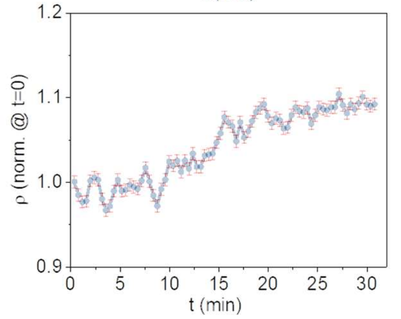


**(a)** High temporal resolution observations of the density dynamics of a single *E. coli* cell. These measurements were taken every 24 seconds. Other than the sampling frequency, the experiment was performed under identical conditions as the data presented in the manuscript (DH5α strain, 1D immobilization in index matched polymers, nutrient supply from top-integrated gels, constant temperature at 37°C, and 63× objective). **(b)** Comparison of the single-cell density quantified via the method presented in this work with alternative segmentation algorithms, including Otsu and Moments. Our method is also compared against a 1D segmentation approach that is independent of conventional thresholding algorithms and, thus, possible errors in area segmentation. While there were slight differences between different methods in the cellular density as determined by each method, all graphs follow the same trend and greatly overlap when normalized at t = 0, with a greater than 99% Pearson correlation coefficient ($p < 0.001$), as displayed in **(c)**. **(d)** Same observations (with the method applied in all experiments), where the blue line denotes the density of the cell and red lines correspond to the error bars reflecting a potential experimental error in plane selection.

### Supplementary Table 1

| sample | test | result |
|---|---|---|
| replicate 1 | Mann-Whitney Test | U = 5657.5; Z = -13.93996; p = 3.62127E-44; |
| replicate 1 | Kolmogorov-Smirnov Test | D = 0.61739; Z = 6.48036; p = 2.85051E-37; |
| replicate 1 | Two sample t Test (Welch Correction) | t = -5.79451; DF = 208; p = 2.51094E-8; |
| replicate 2 | Mann-Whitney Test | U = 21376; Z = -3.31744; p = 9.08477E-4; |
| replicate 2 | Kolmogorov-Smirnov Test | D = 0.17482; Z = 1.6869; p = 0.00576; |
| replicate 2 | Two sample t Test (Welch Correction) | t = -3.5805; DF = 175.03132; p = 4.44038E-4; |
| replicate 3 | Mann-Whitney Test | U = 22010; Z = -4.5231; p = 6.09408E-6; |
| replicate 3 | Kolmogorov-Smirnov Test | D = 0.241; Z = 2.58919; p = 2.25247E-6; |
| replicate 3 | Two sample t Test (Welch Correction) | t = 4.10102; DF = 380.67584; p = 5.03225E-5; |
| all replicates | Mann-Whitney Test | U = 201838; Z = -7.47378; p = 7.79249E-14; |
| all replicates | Kolmogorov-Smirnov Test | D = 0.19379; Z = 3.6008; p = 8.70354E-12; |
| all replicates | Two sample t Test (Welch Correction) | t = 7.6604; DF = 1011.57947; p = 4.32583E-14; |

Various statistical significance tests between growth differentiation ($\gamma_A/\gamma_M$) and cellular dry-density at birth ($\rho_{birth}$). All tests denote that differentiation depends on the density at birth with high statistical significance.
